# Quantifying climate change impacts on plant functional composition and soil nitrogen fixation in Mediterranean grasslands

**DOI:** 10.1101/2022.09.16.508323

**Authors:** Barbara Bomfim, Hilary R. Dawson, Paul B. Reed, Katherine L. Shek, Brendan J. M. Bohannan, Scott D. Bridgham, Lucas C. R. Silva

**Affiliations:** Institute of Ecology and Evolution. University of Oregon, Eugene; Climate and Ecosystem Sciences Division, Lawrence Berkeley National Laboratory; Environmental Studies Program, University of Oregon, Eugene

**Keywords:** Asymbiotic biological nitrogen fixation, disturbance ecology, drought, free-living diazotrophs, temperate grasslands, warming

## Abstract

The projected increase in warming and drought severity (i.e., hotter and drier summers) in the U.S. Pacific Northwest (PNW) may negatively impact grassland plant composition and ecosystem function, with further implications for sustainable land management in the region. To test the vulnerability of Mediterranean grassland function to climate change, we quantified the response of grassland communities to multiannual warming (+2.5°C) and drought (−40% precipitation) by quantifying plant species diversity, legume cover, and biogeochemical controls on and patterns of soil asymbiotic nitrogen fixation (ANF). We hypothesized that the effects of warming on plant functional diversity would increase soil ANF inputs by decreasing legume cover and soil nitrogen availability. Given that asymbiotic N fixers can increase soil organic carbon (C) and nitrogen (N) availability under drought, we hypothesized that the effect of drought on grassland plant cover correlated with increased soil ANF. We surveyed the vegetation and collected composite soil samples from five co-located plots under control (ambient), drought and warming conditions during the fall and spring seasons. In control and drought plots, we quantified the moderator effect of plant composition by comparing low-diversity (unmanipulated plant composition) and high-diversity (manipulated composition) grassland plots. We used a point intercept technique to survey plot-level plant community composition and calculate Shannon’s diversity index and percent cover of legumes (members of Fabaceae according to the Integrated Taxonomic Information System). We measured ANF by incubating collected soils with N-labeled dinitrogen (^15^N_2_), and quantified total soil C, total and available N, available phosphorus (P) and iron (Fe) pools, pH, and soil water holding capacity. Plant species diversity decreased significantly with warming and along the drought severity gradient. ANF response to warming varied by season and site, with rates increasing along the drought severity gradient in the fall but decreasing in the spring. Total soil inorganic N was the strongest predictor of ANF response to warming in the spring but not in the fall. Soil ANF response to drought increased with drought intensity; while soil ANF increased nearly twofold in the southernmost (warm and dry) site, ANF decreased in the northernmost (cool and wet) site. ANF response to drought also varied depending on plant diversity, where low-diversity grasslands had more predictable response to drought than high-diversity grasslands. Soil P availability and pH were the most important variables explaining ANF variability across vegetation types and sites. Our study highlights the importance of using soil-plant-atmosphere interactions to assess grassland ecosystem resilience to drought and warming in the PNW.

## 1 Introduction

Alterations in temperature and precipitation patterns due to climate change (Dai, 2013) are expected to negatively impact Mediterranean grassland ecosystems (Reeves et al., 2014), posing great stress on their vital services and challenging sustainable land use and management. Grassland ecosystems provide key services and livelihood in the U.S. Pacific Northwest (PNW) (Neibergs et al., 2018), where climate change is predicted to reduce summer precipitation, decrease snowpack, and steadily increase air temperatures in the coming century (Neibergs et al., 2018). As a result, grassland ecosystem function, functional diversity and nutrient status are likely to change with decreased water availability and increased temperatures. However, it remains poorly understood how grassland soil-plant interactions will respond to changing temperature and precipitation (Malik & Bouskill, 2022) patterns and how plant functional composition may modulate responses across grassland communities.

Because plant community diversity and nutrient availability are linked, drought and warming impacts on grasslands communities may be highly variable due to complex interactions between the plant community and the environment (Garten et al., 2008). Without climate influence, Busch et al. (2018) showed that nutrient availability had a greater effect than species richness on functional diversity. Without human interference, grassland plants obtain most of their nitrogen (N) via biological N_2_ fixation (BNF), or through plant-associated or free-living diazotrophs (N_2_ fixers), like bacteria or archaea inhabiting the soil (Reed et al., 2011). Asymbiotic N_2_ fixation (ANF) is a process which, unlike mycorrhizal nodes, is more evenly distributed (Sullivan et al., 2014), and accounts for nearly half of the N_2_ fixed via symbiotic BNF rates in temperate grasslands (Reed et al., 2011). Lower BNF rates are predicted with increased temperatures (Garten et al., 2008), and when combined with a decoupling of moisture and N availability (Perez Castro et al., 2020), may alter the provision of atmospheric N needed for photosynthesis.

Under reduced precipitation, plant-soil-microbe interactions can be altered via a reduction in root exudation (Gargallo-Garriga et al., 2018), drought resistance in the microbiome, and a decrease in heterotrophic microbial activity (De Vries et al., 2020); free-living N_2_ fixers can increase soil organic carbon (C) and N availability under drought (Mueller and Bohannan, 2015). However, it remains unclear how chronic climate change affects feedback between plant functional types with symbiotic BNF (e.g., legumes capable of fixing atmospheric N_2_) and free-living BNF across Mediterranean grassland communities (Garten et al., 2008). As a result, it is difficult to predict how the supply of N for grassland plant growth and development (Cavicchioli et al., 2019) will respond to a changing climate.

To test the vulnerability of Mediterranean grassland function to climate change, we investigated the impact of multiannual warming and drought on soil free-living BNF, its major soil biogeochemical controls (Reed et al., 2011; Smercina et al., 2019), and their linkages with plant functional diversity and legume cover. We addressed the following questions: (i) Do warming and drought change the activity of soil diazotrophs across grassland communities? (ii) If so, do changes vary seasonally and relate to plant diversity and legume cover, soil biogeochemistry, and/or meteorology? We tested the hypothesis that warming and drought effects on soil asymbiotic nitrogen fixation (ANF) and biogeochemical controls increase latitudinally along with the Mediterranean drought severity gradient.

We predicted that warming and drought lead to above- and below-ground changes in grassland ecosystems, with the effect of warming on plant functional diversity increasing soil ANF inputs by decreasing legume cover and soil nitrogen availability. Warming would increase soil ANF due to an expected increase in diazotroph activity towards warmer environments (i.e., across the latitudinal gradient and in response to experimental warming at each site). A secondary effect of seasonality on ANF would also be expected as the cold rainy season is expected to decrease diazotroph activity. Given that asymbiotic N fixers can increase soil organic carbon (C) and N availability under drought, we predicted that the effect of drought on grassland plant cover increases soil ANF. To address these hypotheses, we quantified changes in soil free-living BNF rates and carbon, N, phosphorus, iron, pH, moisture and temperature in response to multiannual manipulation of warming (+2.5°C by six 2000-W infrared heaters) and drought (40% reduced precipitation by rain-out shelters) at three grassland sites along a 520 km latitudinal gradient in the U.S. PNW (Reed et al., 2019).

## 2 Materials and Methods

### 2.1 Site description and experimental design

We conducted this study in three experimental grassland sites: Northern (central-western Washington; characterized as cool and wet), Central (central-western Oregon; characterized as warm and wet), and Southern (southwestern Oregon; characterized as warm and dry) (Figure 1; Table 1). Details on the experiment are described elsewhere in greater detail (Reed et al., 2021), so we will only summarize them here. Each site had 25 plots divided into 15 manipulated and 10 unmanipulated plots (vegetation manipulation explained below). The high-diversity plots were further divided into three experimental treatments (5 each of warming, drought, and control). The low-diversity plots were divided into two experimental treatments (5 each of drought and control).

**Figure 1.**
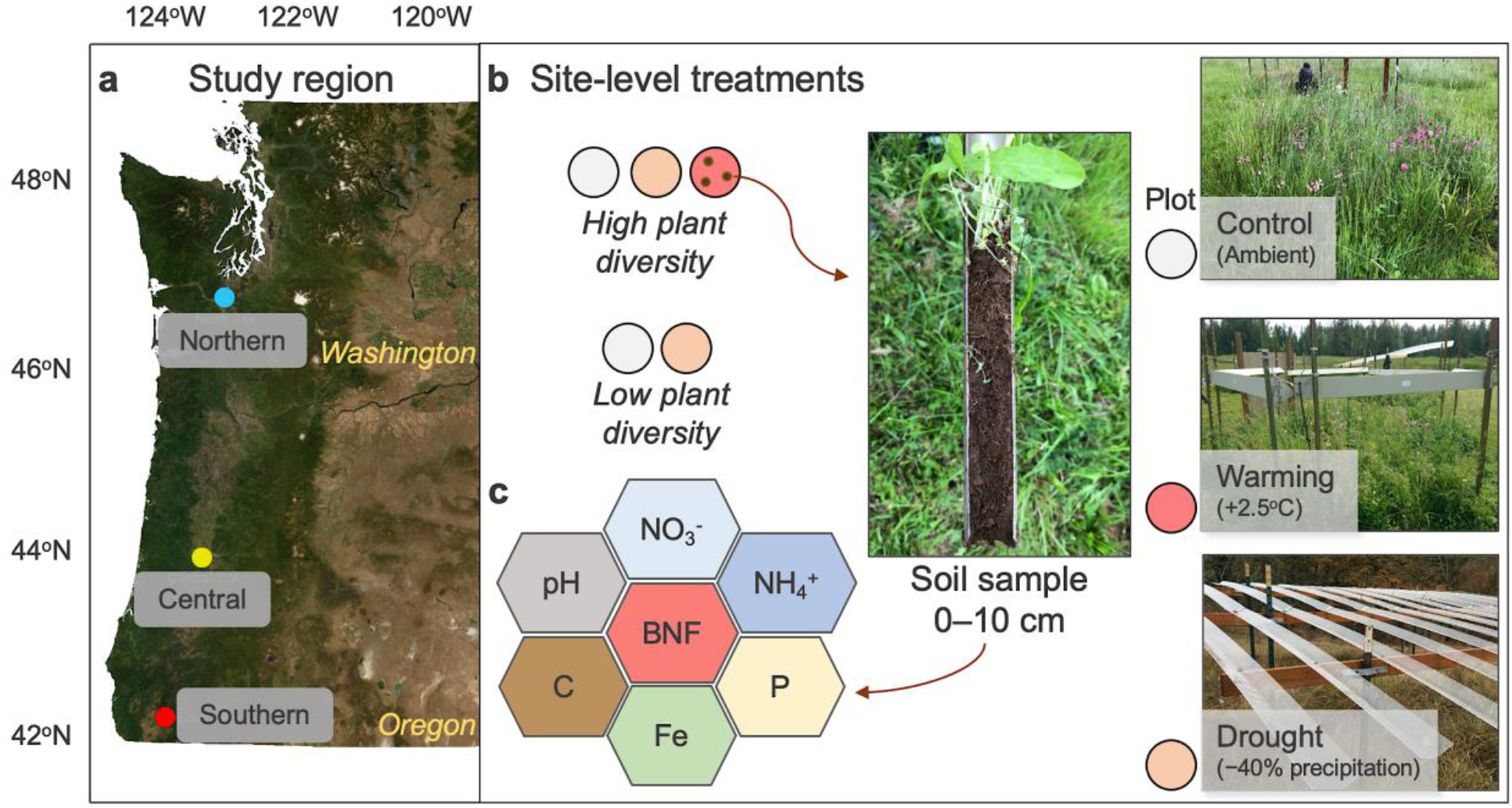
Study sites along the U.S. Pacific Northwest, (a) from southwestern Oregon (Southern) to central-western Oregon (Central) to central-western Washington (Northern). At each site (b), soil samples were collected during the fall and spring seasons from high and low-diversity plots representing three climate treatments: control, drought, and warming (high-diversity plots only). The soils (c) were incubated for ANF measurements, and analyzed for soil biogeochemical variables controlling ANF.

**Table 1.** Soil and environmental variables in each study site along the PNW latitudinal gradient.

| Site |  | Northern |  | Central |  | Southern |  |
| --- | --- | --- | --- | --- | --- | --- | --- |
| Variable | Descriptive statistics | Fall 2018 | Spring 2019 | Fall 2018 | Spring 2019 | Fall 2018 | Spring 2019 |
| Total daily precipitation (mm) | *Mean*<sup>a</sup> | 6.45 | 0.82 | 1.4 | 1.9 | 0.73 | 6.2 |
|  | *Minimum*<sup>b</sup> | 0 | 0 | 0 | 0 | 0 | 0 |
|  | *Maximum*<sup>c</sup> | 36.6 | 4.6 | 14.0 | 14.5 | 8.4 | 66.3 |
| Air temperature (°C) | *Mean* | 4.3 | 12.8 | 11.6 | 13.0 | 10.3 | 10.6 |
|  | *Minimum* | -3.9 | 8.5 | 7.4 | 12.6 | 9.7 | 10.1 |
|  | *Maximum* | 8.4 | 16.9 | 19.9 | 13.5 | 11.0 | 11.2 |
| Total C (g/kg) | *Mean* | 78.5 | 72.5 | 86.7 | 95.9 | 44.6 | 38.7 |
| Total N (g/kg) | *Mean* | 6.4 | 5.8 | 6.5 | 7.2 | 4.1 | 3.5 |
| pH (H<sub>2</sub>O) | *Mean* | 5.8 | 5.8 | 5.7 | 5.8 | 6.4 | 6.3 |
| Soil temperature (°C) | *Mean* | 5.9 | 12.9 | 7.9 | 13.8 | 7.1 | 16.9 |
|  | *Minimum* | 1.0 | 7.5 | 1.1 | 7.8 | 3.7 | 8.9 |
|  | *Maximum* | 8.5 | 17.4 | 16.9 | 22.7 | 10.7 | 32.5 |
| Soil volumetric water content | *Mean* | 0.31 | 0.27 | 0.22 | 0.28 | 0.24 | 0.27 |
|  | *Minimum* | 0.22 | 0.21 | 0.17 | 0.22 | 0.18 | 0.16 |
|  | *Maximum* | 0.46 | 0.37 | 0.35 | 0.36 | 0.35 | 0.40 |

<sup>a</sup> Mean denotes the average among monthly mean soil temperatures measured in control plots at each studied site during the sampling month. <sup>b</sup> Minimum denotes the monthly minimum soil temperature, as described in ‘a’. <sup>c</sup> Maximum denotes the monthly maximum soil temperature, as described in ‘a’. The soil temperature values derive from sensors installed in each studied plot. The precipitation data derived from each site’s meteorological station.

#### Vegetation manipulation

The unmanipulated plots consisted of the vegetation that was already present when the plots were first established. Generally, the unmanipulated plots (hereafter low-diversity) had lower plant diversity than the manipulated plots. The unmanipulated had their species composition unaltered and consisted primarily of the European pasture grasses that dominate at each site. The manipulated plots (hereafter high-diversity) were mowed, raked, sprayed with herbicide, and seeded with a mix of 29 native grasses and forbs. Initial manipulation happened between 2014-2015; afterwards, we annually seeded manipulated plots with 14 native species of grasses and forbs between 2015-2017 (Reed et al. 2020; 2021). Overall, biodiversity of all plots declined over three seasons. Native species showed an exacerbated decline in warming plots (Reed et al., 2021).

#### Climate change experiments

Warming treatment - Warmed plots each had six 2000-W infrared heaters (Kalglo Electronics, Bethlehem, Pennsylvania, USA), which we turned on by 2016, designed to heat the canopy by about 2.5 °C. Previously, we had heated the Southern and Central plots from 2010 to 2012 (Pfeifer-Meister et al., 2013; Reynolds et al., 2015; Wilson et al., 2016b). They were controlled by a dimmer switch using the control plot average to determine heater intensity. We turned off the heaters at all sites in August and September 2017 due to high fire danger. Control plots were ambient temperature plots with ‘dummy heaters’ (wood) installed to mitigate possible shading effects.

Drought treatment – Rain-out shelters covered 40% of the drought plots with clear acrylic shingles (MultiCraft Plastics, Eugene, OR) beginning February 2016 for the manipulated, high-diversity grassland plots and February 2017 for the unmanipulated, low-diversity plots.. The plastic sheeting used absorbed no more than 8% of light transmittance (Yahdjian and Sala, 2002).

### 2.2 Field sampling design

#### Vegetation survey and plant functional data

We used a point intercept technique to survey plot-level plant community composition (see Reed et al. 2021 for further details). In Spring 2017-2019, we measured plant cover at peak standing biomass (mid-May, late-May, and mid-June at the southern, central, and northern sites, respectively) using the point intercept method. Using a 1 m^2^ quadrat in a fixed location within each plot, we dropped 25 equally-spaced pins through the plant canopy to the soil surface. We counted each plant hit on a pin to the species level and multiplied each hit by 4 to standardize to 100. The resulting cover is >100% absolute cover due to the multilayering nature of the canopy. We assigned a cover of 0.4% to species which were present in the quadrat but not hit by a pin. To determine biological N-fixing plant cover, we calculated relative cover as the proportion of absolute cover belonging to members of the plant family Fabaceae.

#### Soil Sampling

In each plot, we used a sterile soil corer to collect three soil samples (0–10 cm depth) combined into one composite sample. Each site was sampled in the fall (November-December 2018) and spring (April-May 2019). The fall sampling included 9 high-diversity and 6 low-diversity plots for each site. All composite soil samples were individually placed into a sterile bag and stored at 4°C during sampling. An additional soil sample was taken from the same plots to determine bulk density (BD). After the sampling was completed in each site, samples were immediately transported to the Soil Plant Atmosphere Research Laboratory at the University of Oregon, Eugene, for the following measurements.

### 2.3 Laboratory measurements

#### Soil physical properties

We determined soil water holding capacity (WHC), gravimetric water content, and BD immediately after sampling. We calculated gravimetric water content (weight of water divided by soil dry weight) by oven-drying ~ 5 g of soil at 100°C for 48 hours (Table S2). We measured bulk density (BD, g cm^-3^) as the total dry weight (oven-dried at 105°C for 48 hours) of the sample divided by the volume of the core sampler (68.71 cm^3^). We determined WHC by placing oven-dried soils in a funnel lined with filter paper over a glass beaker. We added deionized water to the beaker until the soil surface glistened with water. We removed the water in the beaker and left the soil in the funnel to drain. Following draining, we removed the soil and placed it in an oven at 100 °C for 48 h to determine soil moisture content at 100% WHC. We stored all other soils at −20°C for two days while we measured WHC.

#### Soil chemical analyses

After oven-drying for 48 hours, we sieved soil samples to 2 mm, manually ground the soils using a ceramic mortar, and physically and chemically characterized the soils to determine the most important factors known to influence ANF (Reed et al., 2011b). We measured total soil C and N (PDZ Europa ANCA-GSL elemental analyzer), available P and available Fe by Mehlich-3 (M3) extraction (EMBRAPA, 1997; Mehlich, 1984) followed by colorimetric determination (Dominik and Kaupenjohann, 2000; Murphy and Riley, 1962); inorganic N (NH_4_^+^ and NO_3_^-^) using 1M KCl extractions followed by colorimetric determination (Foster, 1995), soil pH in H2O (2:1) and CaCl2 (0.5 M) and soil bulk density (EMBRAPA, 1997).

#### Asymbiotic nitrogen fixation measurements

ANF rates were quantified following Bomfim et al. (2019). We brought all samples to 50% WHC before experimental vials received labeled gas to provide optimal conditions for microbial activity without affecting diffusion of air into samples (Hicks et al., 2003).We incubated three replicates of each soil sample in 12-ml sealed gas vials (Exetainer, Labco, UK) containing 1 g of sample (dry weight) and 2 ml of ^15^N-labeled dinitrogen gas (98 atom%^15^N, catalog number NLM-363-PK, Cambridge Isotope Laboratories), while we incubated three corresponding controls in 12 ml vials containing the same sample (~ 1 g, dry weight) but not receiving any isotopically labeled gas. We incubated all control and labeled vials in a dark incubator (25 °C) for 24 hrs, a period we chose to give ample time for fixation without exhausting the oxygen in the vial. Afterwards, we removed the samples from the vials and measured ANF rates according Hsu and Buckley (2008). We oven-dried all samples at 40 °C and ground in separate ceramic mortars to avoid cross-contamination. N, ^15^N, and C content was determined at the University of California Stable Isotope Facility using a PDZ Europa ANCA-GSL elemental analyzer interfaced to a PDZ Europa 20-20 isotope ratio mass spectrometer (Sercon Ltd., Cheshire, UK). The amount of N fixed (ng N g dry weight^-1^ h^-1^) was calculated in relation to background (control) levels (Table S3).

### 2.4 Statistical analysis

#### Soil ANF and biogeochemistry

We ran all analyses on R version 3.6.2 (R Core Team, 2019). To quantify the effect of each treatment on soil ANF, we first calculated the effect size and the 95% confidence interval for each effect size for each plant diversity treatment and site across both fall and spring seasons. Effect sizes were calculated as the natural log of the response ratio (RR) of soil ANF as following: ln RR = ln (mean ANF treatment plots/mean ANF control plots). Effect sizes are presented in the text as percentages for clarity.

We determined site, plant diversity, climate treatment, and seasonal effects on soil ANF effect sizes and biogeochemical variables by linear mixed-effects models with plot within treatment as a random effect and site, plant diversity, climate treatment, and season as fixed effects. We calculated the conditional (entire model) and marginal (fixed effect) coefficients of determination for the mixed-effect models. Mean effect size values were compared by least-squares means adjusted for Tukey’s HSD test at a 95% significance level.

#### Plant functional diversity

Shannon diversity index was calculated in R using the diversity function in the vegan package (Oksanen et al., 2019) and the point count data. We measured biodiversity using Rao’s Q because of its inclusion of species’ differences in diversity calculations (Botta-Dúkat, 2005). We calculated Rao’s Q in R with the picante package (Kembel et al., 2010) using the community composition data and phylogenetic data derived from the megatree in V.PhyloMaker (Fig. S1; Jin and Hong, 2019; Smith et al., 2018). Relative legume absolute percent cover calculated as legume cover divided by total cover within the quadrat. We then logit-transformed the data using R’s logit function to normalize them as a proportion.

#### Principal component analysis

We conducted a principal component analysis (PCA) to ordinate the plots regarding the main drivers of variation in soil biogeochemistry across sites and climate and diversity treatments. Preliminary PCAs were run using a standardized data matrix including soil ANF, NH_4_, NO_3_+NO_2_, total C/N, Available P, Available Fe, pH, moisture concentration and WHC to select the critical variables for the final PCA. In the preliminary PCA, low (< 1.0) eigenvalues (Peña-Claros et al., 2012) were found for all parameters except total C/N, available P, available Fe, NH_4_, moisture and pH, which were thus included in the final PCA.

## 3 Results

### 3.1 Seasonal climate change effects on soil ANF across sites and plant diversity treatments

We detected ANF activity in 78% (94 out of 120) of the plots sampled across, sites, climate and plant diversity treatments during the fall and spring seasons. Average ANF rates varied substantially across measurements, ranging between 0.41 ng N g dry weight^-1^ h^-1^ in the warming treatment of the southern site and 5.54 ng N g dry weight^-1^ h^-1^ in the control plots of the central site (Figure S1). ANF effect sizes ranged from −42.3% (SD 36.8%) in the warming treatment of the southern site in the Spring and 25.9% (SD 10.5%) in the drought treatment of the southern site in the Fall (Figure 2).

**Figure 2.**
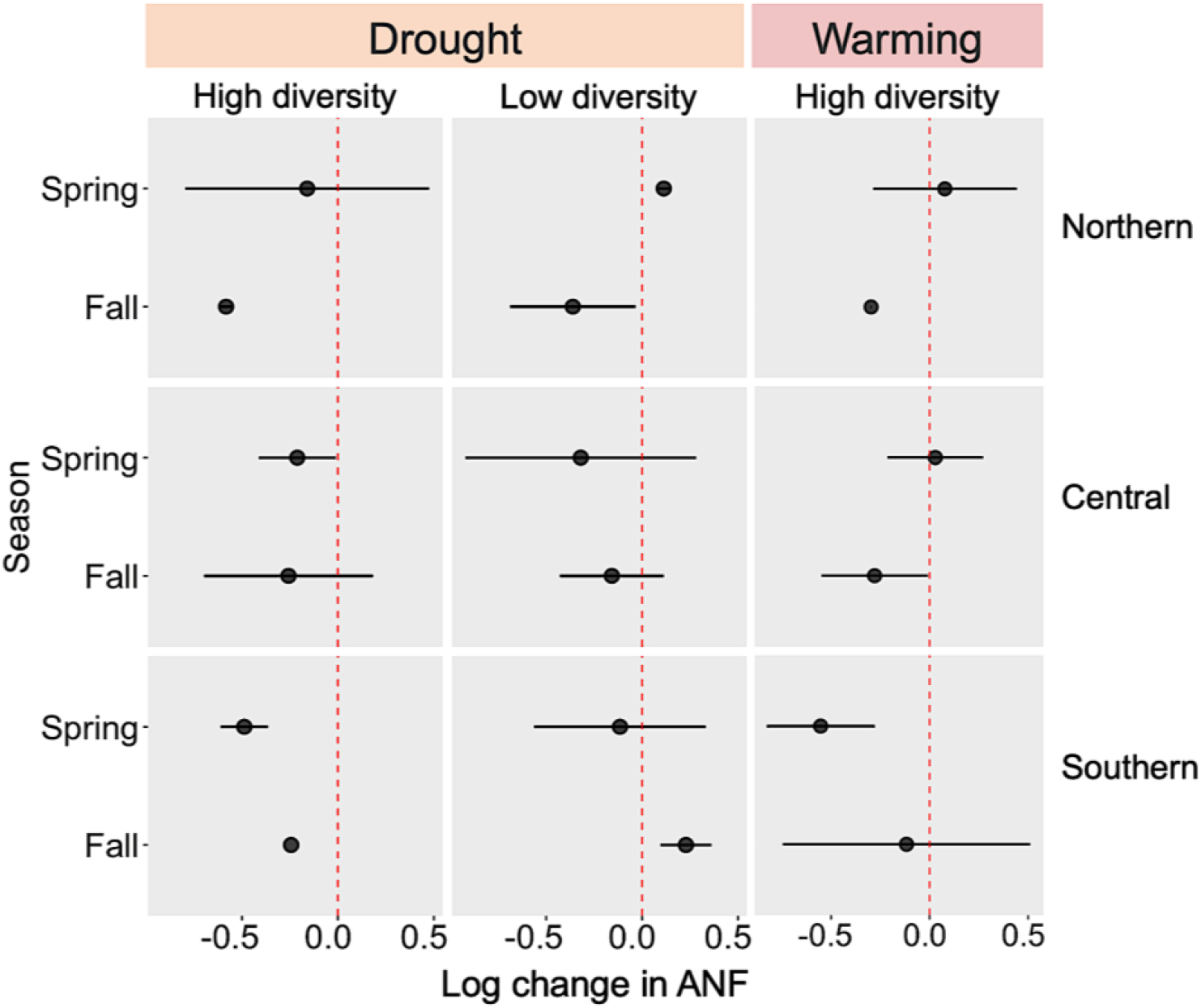
Drought and warming effects on asymbiotic nitrogen fixation - mean log10 (ANF treatment/control) with 95% confidence interval – in high and low plant diversity plots established in grasslands across three study sites located along a > 500km latitudinal gradient in the Pacific Northwest. The vertical dashed lines indicate the control conditions. Points are means and horizontal lines indicate the 95% confidence interval.

Soil ANF response to drought increased along the drought intensity gradient in the Fall season (Figure 2). While ANF rates increased nearly twofold in the southernmost site, significant decrease in ANF was verified in the northernmost site. ANF response to drought also varied depending on plant diversity, where low-diversity grasslands had more predictable response to drought than high-diversity grasslands. For instance, ANF in southern high-diversity grasslands was suppressed but no change was verified in central high-diversity grasslands.

Among drought plots, we found a significant effect of plant diversity (as a categorical variable) on log change in ANF (Kruskal-Wallis chi-squared = 3.81, df = 1, p-value = 0.048). Among warming plots, no effect of site or season on log change in ANF. ANF response to warming varied by season and site, where rates increased with the drought severity gradient in the fall but decreased during the spring.

### 3.2 Soil biogeochemical responses to warming and drought and controls on ANF

Total soil C and N contents were generally higher in high-diversity grasslands whereas warming treatment had no significant effect across sites. Total soil inorganic nitrogen availability (sum of nitrate, nitrite and ammonium) was the strongest predictor of ANF response to warming in the spring but not in the fall. Soil P availability, also affected by drought, and pH were the most important variables explaining ANF variability across vegetation types and sites.

The PCA indicated that moisture grouped with total soil C and N in opposed quadrants to pH (Figure 3). On the other hand, ammonium co-varied with available P and Fe, all orthogonally to the soil C, N, moisture and pH axis.

**Figure 3.**
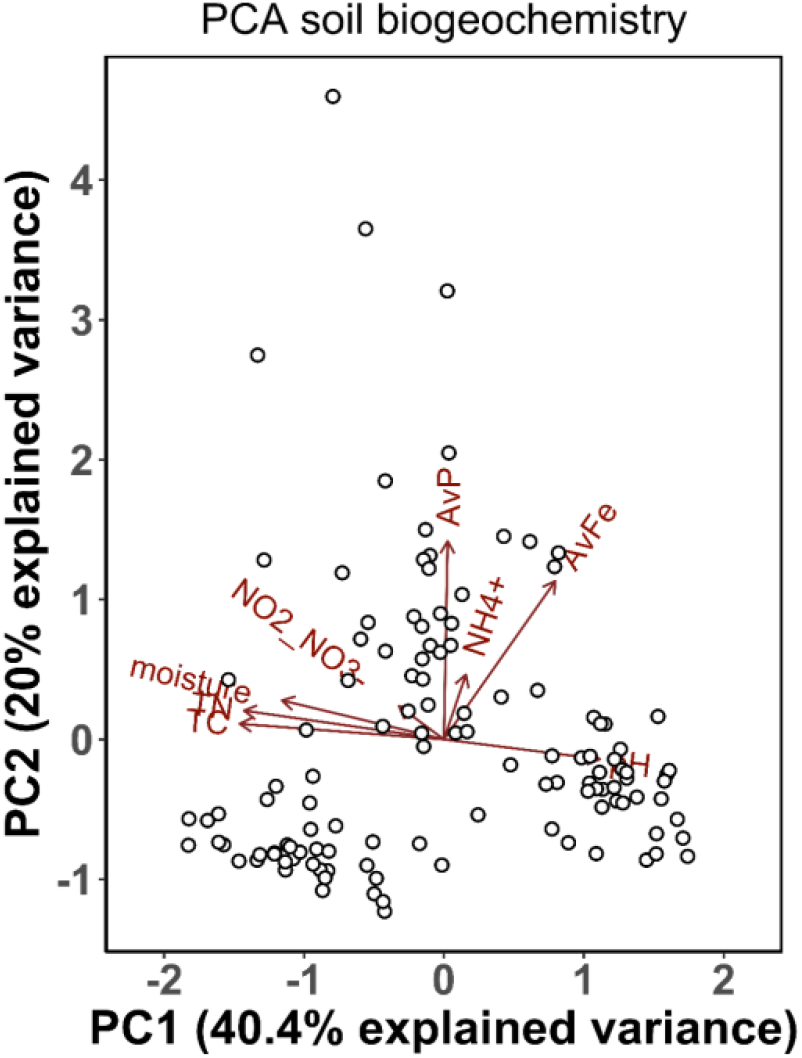
Principal component analysis (PCA) of soil ANF (ng N g dry weight^-1^ h^-1^), NH_4_ (μg g dry weight^-1^) NO_3_+NO_2_ (μg g dry weight^-1^), total C/N (mass-based), Available P (μg g dry weight^-1^), Available Fe (μg g dry weight^-1^), pH and moisture concentration (%). Each point represents the plot averages of these variables, in a total of 75 plots representing all sites and seasons. PC1 (x-axis) explained 40.4%, and PC2 (y-axis) explained 20% of the total variance in the soil biogeochemistry data.

### 3.3 Plant functional diversity and Legume cover

High diversity plots were twice as diverse as low diversity plots (Figure 4a). There was a significant interaction between site and plot diversity (Table S4). The northern and central high diversity plots as the most diverse (Rao’s Q = 72.3 ± 22.9 and 71.7 ± 14.3, respectively), while northern low diversity plots were the least diverse (Rao’s Q = 10.2 ± 3.20). Warming plots were nearly twice as diverse compared to control and drought plots, which did not differ (Figure 4a; Table S4).

**Figure 4.**
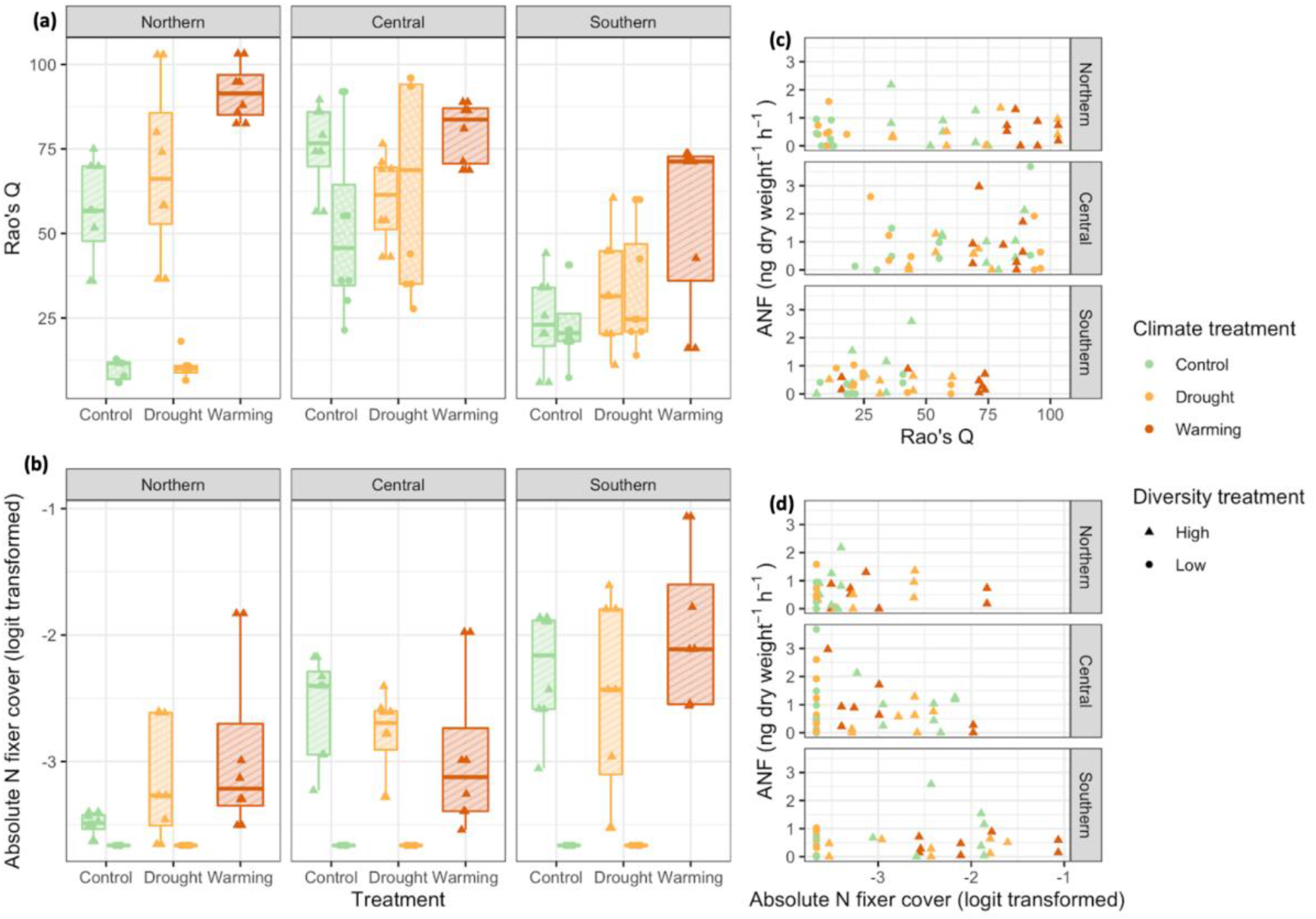
Treatment effect on Rao’s Q, biological N-fixing plant cover, and correlation with soil ANF (in ng N g dry weight^-1^ h^-1^).

All sites were significantly different in terms of N-fixing plant cover (Figure 4b; Table S4). Low diversity plots did not have any N-fixing plant cover (Figure 4b). In high diversity plots, the northern site had the lowest N-fixing plant cover (1.29% ± 2.76) and the southern site had the highest (5.28% ± 6.53). Climate treatment and season of sampling did not change N-fixing plant cover, which increased significantly from 3.1% in the northern to 9.2% in the southernmost site. We found no correlation between N-fixing plant cover or Rao’s Q and ANF (Figure 4c-d).

## 4 Discussion

Understanding how climate modulates the relationship between nutrient availability and plant functional diversity is important because grasslands with higher plant species diversity show higher inter-specific resource transfer and community resilience. To test the vulnerability of Mediterranean grassland function to climate change, we quantified the response of grassland communities to multiannual warming and drought by quantifying plant species diversity, legume cover, and biogeochemical controls on and patterns of soil asymbiotic nitrogen fixation (ANF). Our results partially support our hypothesis that the effects of warming on plant functional diversity would increase soil ANF inputs by decreasing legume cover and soil nitrogen availability.

### 4.1 Drought and warming effects on soil ANF and biogeochemistry

Because N_2_ fixation is an enzymatic process, we expected to see a rate increase that corresponds with soil temperature until the enzyme denatures (Reed et al., 2011a). Based on our findings, low-diversity grasslands in the northern site may be those most severely impacted by increased climate change induced drought stress. This is expected because drought can impart changes in plant and microbial communities that can lead to changes in plant litter chemistry and the chemical composition of microbial necromass, both of which are key resources for soil microbes (Malik & Bouskill, 2022).

The dependence of reaction rates on temperature, including the rates of enzymatic reactions, is described by the Arrhenius equation (Davidson and Janssens, 2006). According to this equation, the fractional increase in reaction rate is less for a temperature increase of one degree Celsius at higher temperatures than at lower temperatures. This suggests that the response of soil microbial respiration to temperature changes should be less pronounced in tropical climates than it is in temperate or boreal climates.

Little is known about ANF responses to rhizosphere conditions. Controls on ANF may include C, N, O, and P availability as well as pH, temperature, and soil moisture and temperature but more research is needed (Smercina et al., 2019). Lower sampling soil temperature was correlated with reduced N fixation in cyanobacteria when incubated under constant temperature (Liengen and Olsen, 1997). Similarly, increased long-term incubation temperature significantly increased microbial respiration (Zogg et al., 1997). One might expect to see N-rich compatible solutes (e.g. proline, ectoine) dominate the drought response in N rich soils versus C-rich compatible solutes (e.g trehalose) in N-poor soils. Those will have different costs to the microbe so that might be one explanation for the interaction between nutrient status and drought adaptation of microbes.

### 4.2 Plant functional diversity across treatments

Understanding changes in plant functional diversity and grassland function due to climate change matters regionally because rangeland, pasture, and grazed federal lands represent the largest land areas in the PNW (Neibergs et al., 2018). A previous study in the same sites showed that warming increased the cover of introduced annual species, causing subse-quent declines in other functional groups and diversity (Reed et al., 2021). The authors reported that competition for moisture and light or space, rather than nitro-gen, were critical mechanisms of community change in these seasonally water-limited Mediterranean grasslands. Their findings corroborate their previous research suggesting that future climate change will alter plant community composition and decrease diversity in PNW grasslands.

Climate change impacts on grassland N cycling needs to be better predicted because it is expected that severe droughts and exotic plants may produce a larger, more vulnerable pool of N that is prone to losses while providing a competitive advantage to promote exotic growth in grassland N-limited ecosystems (Pérez Castro et al., 2020). More intense and prolonged droughts due to climate change are expected to promote open N cycling in semi-arid shrublands via sustained N processing during periods of low plant uptake (Evans and Burke, 2012). In this context, herbaceous plants capable of fixing N can modulate the impacts of climate change on soil N.

However, there has been little attention given to N-fixing grassland species in the context of climate change. The most well-known N-fixing plant family is Fabaceae (legumes); however, other families in the Pacific Northwest have some species with N-fixing symbiosis including Betulaceae, Myrticaceae, Rosaceae, and Rhamnaceae (Tedersoo et al., 2018). Researchers have also observed associations between Poaceae (grasses) and N_2_-fixing bacteria, particularly in commercial crops such as sugar cane, wheat, and rice (Pankievicz et al., 2015). There are fewer data available on wild species and the commonality of such associations. Legumes can reduce respiration and microbial C:N in grasslands (compared to other forbs and grasses) and may lead to phosphorus limitation (Strecker et al., 2015).

## 5 Conclusions

We found that the effects of warming and drought on soil ANF and its biogeochemical controls varie considerably across the drought intensity gradient in the U.S. PNW. This suggests that extrapolating site-level results to regional scale may not lead to accurate predictions of how grassland ecosystems will respond and adapt to future climate change scenarios. This study highlights the importance of using soil-plant-atmosphere interactions to assess grassland resilience to climate change in the PNW.

## Acknowledgements

Siskiyou Field Institute, The Nature Conservancy, and Capitol Land Trust provided sites for this experiment, Laurel Pfeifer-Meister, Bitty Roy, Bart Johnson, Graham Bailes, Aaron Nelson, and Matthew Krna contributed to the experimental design, and numerous others assisted with the HOPS project. This experiment was funded by National Science Foundation Macrosystems Biology grant #1340847, Plant Biotic Interactions grant #1758947, and Convergence Accelerator Pilot grant #1939511.

## Notes

### Competing Interest Statement

The authors have declared no competing interest.

### Summary of Updates

Author names updated; Minor update in the Abstract and keywords.

