## Supplementary Material for "Quantifying climate change impacts on plant functional composition and soil nitrogen fixation in Mediterranean grasslands"

### Supplementary Materials

**Table S1** Geographical coordinates of the plots sampled across the three sites where the climate change experiment was conducted.

| Site ID | Plot ID | Plant diversity | Treatment | Latitude | Longitude |
|---|---|---|---|---|---|
| Southern | 1 | High | Drought | 42°16'41.46" | 123°38'31.58" |
| Southern | 2 | High | Control | 42°16'41.49" | 123°38'31.85" |
| Southern | 4 | High | Drought | 42°16'41.53" | 123°38'32.36" |
| Southern | 5 | High | Warming | 42°16'41.55" | 123°38'32.63" |
| Southern | 7 | High | Drought | 42°16'41.28" | 123°38'31.87" |
| Southern | 8 | High | Warming | 42°16'41.31" | 123°38'32.14" |
| Southern | 10 | High | Control | 42°16'41.35" | 123°38'32.67" |
| Southern | 11 | High | Warming | 42°16'41.06" | 123°38'31.66" |
| Southern | 13 | High | Control | 42°16'41.10" | 123°38'32.18" |
| Southern | 14 | High | Warming | 42°16'41.15" | 123°38'32.69" |
| Southern | 16 | High | Control | 42°16'40.85" | 123°38'31.68" |
| Southern | 17 | High | Warming | 42°16'40.86" | 123°38'31.95" |
| Southern | 18 | High | Drought | 42°16'40.88" | 123°38'32.20" |
| Southern | 19 | High | Control | 42°16'40.91" | 123°38'32.47" |
| Central | 21 | High | Drought | 44° 1'34.68" | 123°10'56.48" |
| Central | 22 | High | Control | 44° 1'34.47" | 123°10'56.45" |
| Central | 23 | High | Warming | 44° 1'34.26" | 123°10'56.47" |
| Central | 25 | High | Drought | 44° 1'33.85" | 123°10'56.48" |
| Central | 29 | High | Control | 44° 1'33.85" | 123°10'56.21" |
| Central | 30 | High | Drought | 44° 1'34.49" | 123°10'55.94" |
| Central | 32 | High | Warming | 44° 1'34.06" | 123°10'55.96" |
| Central | 34 | High | Control | 44° 1'34.49" | 123°10'55.67" |
| Central | 35 | High | Warming | 44° 1'34.27" | 123°10'55.66" |
| Northern | 42 | High | Warming | 46°51'50.97" | 122°57'32.27" |
| Northern | 47 | High | Drought | 46°51'51.15" | 122°57'32.00" |
| Northern | 48 | High | Control | 46°51'51.16" | 122°57'31.71" |
| Northern | 50 | High | Control | 46°51'51.36" | 122°57'32.29" |
| Northern | 51 | High | Warming | 46°51'51.37" | 122°57'31.99" |
| Northern | 56 | High | Drought | 46°51'51.55" | 122°57'31.72" |
| Northern | 58 | High | Drought | 46°51'51.74" | 122°57'32.29" |
| Northern | 59 | High | Control | 46°51'51.75" | 122°57'32.00" |
| Northern | 60 | High | Warming | 46°51'51.75" | 122°57'31.72" |
| Northern | 121 | Low | Control | 46°51'50.12" | 122°57'32.62" |

| | | | | | |
|---|---|---|---|---|---|
| Northern | 122 | Low | Drought | 46°51'50.15" | 122°57'32.87" |
| Northern | 123 | Low | Control | 46°51'50.16" | 122°57'33.09" |
| Northern | 126 | Low | Drought | 46°51'50.00" | 122°57'33.13" |
| Northern | 127 | Low | Drought | 46°51'49.83" | 122°57'32.70" |
| Northern | 128 | Low | Control | 46°51'49.83" | 122°57'32.93" |

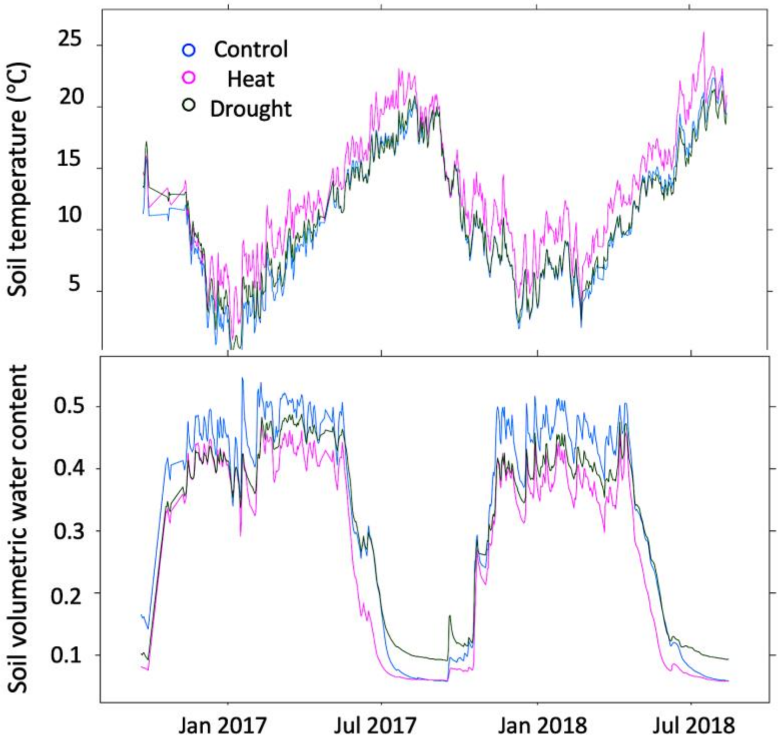


**Figure S1** Mean soil volumetric water content and temperature (in degrees Celsius) measured in control (blue), heat (purple) and drought (green) high-diversity plots between late Summer 2017 and Summer 2018.

**Table S2.** Soil physical variables measured across sites, plots, seasons, and plant diversity and climate treatments. BD denotes bulk density and WHC water holding capacity.

| **Site** | **Plot ID** | **Season** | **Plant diversity** | **Treatment** | **BD ($g/cm^3$)** | **Gravimetric water content (%)** | **WHC (mg/g)** |
|---|---|---|---|---|---|---|---|
| Northern | 41 | Spring | High | Control | 0.44 | 22.45 | 810.51 |
| | 42 | Spring | High | Warming | 0.60 | 21.39 | 460.52 |
| | 42 | Fall | High | Warming | 0.60 | 60.00 | 402.08 |
| | 44 | Spring | High | Drought | 0.63 | 22.94 | 818.62 |
| | 45 | Spring | High | Warming | 0.58 | 28.47 | 334.58 |
| | 47 | Spring | High | Drought | 0.27 | 25.06 | 781.90 |
| | 47 | Fall | High | Drought | 0.27 | 44.99 | 611.34 |
| | 48 | Spring | High | Control | 0.41 | 29.11 | 709.44 |
| | 48 | Fall | High | Control | 0.41 | 39.95 | 715.03 |
| | 49 | Spring | High | Drought | 0.80 | 25.94 | 765.90 |
| | 50 | Spring | High | Control | 0.43 | 25.72 | 773.23 |
| | 50 | Fall | High | Control | 0.43 | 41.91 | 608.41 |
| | 51 | Spring | High | Warming | 0.47 | 24.12 | 682.21 |
| | 51 | Fall | High | Warming | 0.47 | 38.78 | 628.64 |

| | | | | | | | |
|---|---|---|---|---|---|---|---|
| | 53 | Spring | High | Control | 0.64 | 28.25 | 726.19 |
| | 54 | Spring | High | Warming | 0.84 | 27.39 | 352.72 |
| | 56 | Fall | High | Drought | 0.46 | 38.45 | 494.66 |
| | 56 | Spring | High | Drought | 0.46 | 23.76 | 566.85 |
| | 58 | Fall | High | Drought | 0.82 | 30.31 | 622.33 |
| | 58 | Spring | High | Drought | 0.82 | 24.30 | 434.00 |
| | 59 | Fall | High | Control | 0.47 | 40.30 | 515.02 |
| | 59 | Spring | High | Control | 0.47 | 28.31 | 745.03 |
| | 60 | Fall | High | Warming | 0.45 | 39.04 | 676.96 |
| | 60 | Spring | High | Warming | 0.45 | 27.18 | 802.84 |
| | 121 | Spring | Low | Control | 0.63 | 28.60 | 560.12 |
| | 121 | Fall | Low | Control | 0.63 | 34.20 | 542.49 |
| | 122 | Spring | Low | Drought | 0.18 | 23.79 | 723.59 |
| | 122 | Fall | Low | Drought | 0.18 | 44.58 | 1439.96 |
| | 123 | Spring | Low | Control | 0.19 | 40.79 | 299.46 |
| | 123 | Fall | Low | Control | 0.19 | 46.25 | 491.39 |
| | 124 | Spring | Low | Drought | 0.53 | 18.06 | 809.63 |
| Northern | 125 | Spring | Low | Control | 0.61 | 25.43 | 658.95 |
| | 126 | Spring | Low | Drought | 0.44 | 35.36 | 479.66 |
| | 126 | Fall | Low | Drought | 0.44 | 41.45 | 485.87 |
| | 127 | Spring | Low | Drought | 0.77 | 29.73 | 647.12 |
| | 127 | Fall | Low | Drought | 0.77 | 31.36 | 681.88 |
| | 128 | Spring | Low | Control | 0.35 | 45.32 | 558.76 |
| | 128 | Fall | Low | Control | 0.35 | 44.42 | 265.25 |
| | 129 | Spring | Low | Control | 0.55 | 29.84 | 564.68 |
| | 130 | Spring | Low | Drought | 0.57 | 26.67 | 676.31 |
| | 21 | Fall | High | Drought | 0.35 | 19.56 | 900.26 |
| | 21 | Spring | High | Drought | 0.35 | 36.81 | 657.12 |
| | 22 | Fall | High | Control | 0.50 | 28.04 | 884.53 |
| | 22 | Spring | High | Control | 0.50 | 37.40 | 494.92 |
| | 23 | Fall | High | Warming | 0.57 | 28.37 | 838.03 |
| | 23 | Spring | High | Warming | 0.57 | 29.11 | 441.95 |
| | 25 | Fall | High | Drought | 0.58 | 24.44 | 961.04 |
| | 25 | Spring | High | Drought | 0.58 | 36.76 | 637.51 |
| | 26 | Spring | High | Warming | 0.10 | 30.63 | 726.61 |
| | 28 | Spring | High | Drought | 0.11 | 43.80 | 857.62 |
| | 29 | Fall | High | Control | 0.57 | 25.86 | 1073.90 |
| Central | 29 | Spring | High | Control | 0.57 | 47.24 | 725.07 |
| | 30 | Fall | High | Drought | 0.55 | 26.67 | 1005.18 |
| | 30 | Spring | High | Drought | 0.55 | 41.98 | 641.55 |
| | 31 | Spring | High | Control | 0.35 | 52.51 | 693.04 |
| | 32 | Fall | High | Warming | 0.55 | 29.66 | 1185.19 |
| | 32 | Spring | High | Warming | 0.55 | 30.69 | 763.45 |
| | 34 | Fall | High | Control | 0.65 | 32.54 | 1183.62 |
| | 34 | Spring | High | Control | 0.65 | 38.84 | 701.26 |
| | 35 | Fall | High | Warming | 0.47 | 33.94 | 1159.42 |
| | 35 | Spring | High | Warming | 0.47 | 28.40 | 645.59 |
| | 38 | Spring | High | Drought | 0.45 | 38.72 | 714.91 |
| | 39 | Spring | High | Control | 0.41 | 34.94 | 824.50 |
| | 40 | Spring | High | Warming | 0.60 | 37.92 | 676.43 |
| | 111 | Fall | Low | Control | 0.45 | 30.00 | 964.04 |
| | 111 | Spring | Low | Control | 0.45 | 47.59 | 535.47 |
| Central | 112 | Fall | Low | Drought | 0.46 | 31.09 | 1162.79 |
| | 112 | Spring | Low | Drought | 0.46 | 47.70 | 711.63 |
| | 113 | Spring | Low | Control | 0.30 | 39.62 | 622.33 |

| | | | | | | | |
|---|---|---|---|---|---|---|---|
| | 114 | Spring | Low | Drought | 0.24 | 44.02 | 801.17 |
| | 115 | Fall | Low | Control | 0.28 | 36.45 | 1494.32 |
| | 115 | Spring | Low | Control | 0.28 | 34.18 | 920.86 |
| | 116 | Spring | Low | Control | 0.36 | 39.82 | 909.85 |
| | 117 | Fall | Low | Drought | 0.41 | 33.25 | 1145.58 |
| | 117 | Spring | Low | Drought | 0.41 | 42.41 | 647.38 |
| | 118 | Spring | Low | Drought | 0.21 | 42.48 | 793.54 |
| | 119 | Fall | Low | Control | 0.42 | 36.74 | 1451.87 |
| | 119 | Spring | Low | Control | 0.42 | 40.45 | 871.06 |
| | 120 | Fall | Low | Drought | 0.36 | 35.74 | 1339.57 |
| | 120 | Spring | Low | Drought | 0.36 | 45.10 | 783.50 |
| | 1 | Fall | High | Drought | 0.92 | 29.11 | 560.34 |
| | 1 | Spring | High | Drought | 0.92 | 14.24 | 394.25 |
| | 2 | Fall | High | Control | 0.85 | 29.15 | 753.47 |
| | 2 | Spring | High | Control | 0.85 | 10.35 | 638.09 |
| | 4 | Spring | High | Drought | 0.99 | 11.51 | 531.65 |
| | 5 | Fall | High | Warming | 1.21 | 25.20 | 687.50 |
| | 5 | Spring | High | Warming | 1.21 | 11.29 | 560.19 |
| | 7 | Fall | High | Drought | 1.09 | 26.27 | 650.64 |
| | 7 | Spring | High | Drought | 1.09 | 11.40 | 521.99 |
| | 8 | Fall | High | Warming | 1.35 | 21.67 | 670.42 |
| | 8 | Spring | High | Warming | 1.35 | 10.41 | 554.25 |
| Southern | 10 | Fall | High | Control | 0.86 | 28.01 | 737.83 |
| | 10 | Spring | High | Control | 0.86 | 11.89 | 602.94 |
| | 11 | Spring | High | Warming | 1.00 | 18.06 | 428.16 |
| | 13 | Spring | High | Control | 1.09 | 15.96 | 408.99 |
| | 14 | Spring | High | Warming | 0.82 | 14.89 | 359.60 |
| | 15 | Spring | High | Drought | 1.20 | 15.50 | 415.40 |
| | 16 | Spring | High | Control | 0.93 | 17.49 | 451.13 |
| | 17 | Fall | High | Warming | 1.09 | 26.89 | 795.62 |
| | 17 | Spring | High | Warming | 1.09 | 13.93 | 633.79 |
| | 18 | Fall | High | Drought | 1.09 | 25.28 | 646.50 |
| | 18 | Spring | High | Drought | 1.09 | 11.67 | 514.37 |
| | 19 | Fall | High | Control | 0.89 | 26.77 | 514.71 |
| | 19 | Spring | High | Control | 0.89 | 15.13 | 336.47 |
| | 101 | Fall | Low | Drought | 1.04 | 28.52 | 794.57 |
| | 101 | Spring | Low | Drought | 1.04 | 14.45 | 625.74 |
| | 102 | Fall | Low | Control | 1.03 | 32.19 | 935.48 |
| | 102 | Spring | Low | Control | 1.03 | 15.29 | 755.05 |
| | 103 | Fall | Low | Control | 1.07 | 31.86 | 1152.42 |
| | 103 | Spring | Low | Control | 1.07 | 16.89 | 949.17 |
| | 104 | Spring | Low | Control | 0.63 | 15.66 | 637.18 |
| Southern | 105 | Spring | Low | Drought | 1.01 | 16.87 | 638.91 |
| | 106 | Fall | Low | Drought | 0.88 | 28.57 | 981.19 |
| | 106 | Spring | Low | Drought | 0.88 | 14.89 | 806.23 |
| | 107 | Spring | Low | Control | 0.73 | 13.16 | 570.57 |
| | 108 | Fall | Low | Control | 0.89 | 29.71 | 1064.06 |
| | 108 | Spring | Low | Control | 0.89 | 16.55 | 865.70 |
| | 109 | Fall | Low | Drought | 0.89 | 31.44 | 1000.00 |
| | 109 | Spring | Low | Drought | 0.89 | 13.47 | 844.35 |
| | 110 | Spring | Low | Drought | 0.65 | 13.50 | 639.48 |

**Table S3.** Soil asymbiotic nitrogen fixation rates (ANF, ng N g dry weight$^{-1}$ h$^{-1}$) calculated for all samples collected during the Fall and Spring seasons from all three site and treatments. Effect sizes are the $\log_{10}$ transformation of the ratio of ANF in a sample from a climate treatment to the mean ANF in the control samples for the same site, plant diversity, and season. Fixation activity indicates whether ANF rate was detected, where N indicates 'no fixation' and Y indicates fixation was detected and calculated.

| Site ID | Plot ID | Season | Plant diversity | Climate treatment | Fixation | ANF (ng N/g/h) | Effect size [a] $\log_{10}$ (climate treatment/control) |
|---|---|---|---|---|---|---|---|
| Northern | 41 | Spring | High | Control | N | 0.001 | - |
| Northern | 42 | Fall | High | Warming | N | 0.001 | - |
| Northern | 42 | Spring | High | Warming | Y | 0.88 | 0.264 |
| Northern | 44 | Spring | High | Drought | Y | 0.02 | -1.379 |
| Northern | 45 | Spring | High | Warming | N | 0.001 | - |
| Northern | 47 | Fall | High | Drought | Y | 0.362 | -0.602 |
| Northern | 47 | Spring | High | Drought | Y | 0.301 | -0.202 |
| Northern | 48 | Fall | High | Control | Y | 0.904 | - |
| Northern | 48 | Spring | High | Control | Y | 0.503 | - |
| Northern | 49 | Spring | High | Drought | Y | 1.357 | 0.452 |
| Northern | 50 | Fall | High | Control | Y | 1.26 | - |
| Northern | 50 | Spring | High | Control | Y | 0.127 | - |
| Northern | 51 | Fall | High | Warming | Y | 0.728 | -0.299 |
| Northern | 51 | Spring | High | Warming | Y | 0.52 | 0.036 |
| Northern | 53 | Spring | High | Control | N | 0.001 | - |
| Northern | 54 | Spring | High | Warming | Y | 1.3 | 0.434 |
| Northern | 56 | Fall | High | Drought | N | 0.001 | - |
| Northern | 56 | Spring | High | Drought | Y | 0.512 | 0.029 |
| Northern | 58 | Fall | High | Drought | Y | 0.395 | -0.564 |
| Northern | 58 | Spring | High | Drought | Y | 0.953 | 0.299 |
| Northern | 59 | Fall | High | Control | Y | 2.18 | - |
| Northern | 59 | Spring | High | Control | Y | 0.807 | - |
| Northern | 60 | Fall | High | Warming | Y | 0.7339 | -0.295 |
| Northern | 60 | Spring | High | Warming | Y | 0.18 | -0.425 |
| Northern | 121 | Fall | Low | Control | Y | 0.925 | -0.427 |
| Northern | 121 | Spring | Low | Control | N | 0.001 | - |
| Northern | 122 | Fall | Low | Drought | Y | 0.732 | -0.528 |
| Northern | 122 | Spring | Low | Drought | Y | 0.413 | 0.085 |
| Northern | 123 | Fall | Low | Control | Y | 0.948 | - |
| Northern | 123 | Spring | Low | Control | Y | 0.443 | - |
| Northern | 124 | Spring | Low | Drought | N | 0.001 | - |
| Northern | 125 | Spring | Low | Control | Y | 0.236 | -0.158 |
| Northern | 126 | Fall | Low | Drought | N | 0.001 | - |
| Northern | 126 | Spring | Low | Drought | Y | 0.448 | 0.12 |
| Northern | 127 | Fall | Low | Drought | Y | 1.582 | -0.194 |
| Northern | 127 | Spring | Low | Drought | Y | 0.498 | 0.166 |
| Northern | 128 | Fall | Low | Control | Y | 5.54 | 0.351 |
| Northern | 128 | Spring | Low | Control | N | 0.001 | - |
| Northern | 129 | Spring | Low | Control | N | 0.001 | - |
| Northern | 130 | Spring | Low | Drought | Y | 0.41 | 0.082 |
| Central | 21 | Fall | High | Drought | Y | 0.627 | -0.013 |
| Central | 21 | Spring | High | Drought | Y | 1.284 | -0.019 |
| Central | 22 | Fall | High | Control | Y | 0.434 | - |
| Central | 22 | Spring | High | Control | Y | 1.035 | - |
| Central | 23 | Fall | High | Warming | Y | 0.6342 | -0.008 |

| | | | | | | | |
|---|---|---|---|---|---|---|---|
| Central | 23 | Spring | High | Warming | Y | 1.71 | 0.106 |
| Central | 25 | Fall | High | Drought | Y | 0.127 | -0.707 |
| Central | 25 | Spring | High | Drought | N | 0.001 | -3.127 |
| Central | 26 | Spring | High | Warming | Y | 0.89 | -0.178 |
| Central | 28 | Spring | High | Drought | Y | 0.754 | -0.25 |
| Central | 29 | Fall | High | Control | Y | 1.262 | - |
| Central | 29 | Spring | High | Control | Y | 1.189 | - |
| Central | 30 | Fall | High | Drought | Y | 0.573 | -0.053 |
| Central | 30 | Spring | High | Drought | Y | 0.576 | -0.367 |
| Central | 31 | Spring | High | Control | Y | 2.13 | - |
| Central | 32 | Fall | High | Warming | Y | 0.2258 | -0.457 |
| Central | 32 | Spring | High | Warming | Y | 0.93 | -0.159 |
| Central | 34 | Fall | High | Control | Y | 0.244 | - |
| Central | 34 | Spring | High | Control | Y | 1.006 | - |
| Central | 35 | Fall | High | Warming | Y | 0.2745 | -0.372 |
| Central | 35 | Spring | High | Warming | N | 0.001 | - |
| Central | 38 | Spring | High | Drought | N | 0.001 | - |
| Central | 39 | Spring | High | Control | N | 0.001 | - |
| Central | 40 | Spring | High | Warming | Y | 2.97 | 0.346 |
| Central | 111 | Fall | Low | Control | Y | 0.522 | - |
| Central | 111 | Spring | Low | Control | Y | 3.685 | - |
| Central | 112 | Fall | Low | Drought | N | 0.001 | - |
| Central | 112 | Spring | Low | Drought | Y | 1.92 | 0.129 |
| Central | 113 | Spring | Low | Control | Y | 0.133 | - |
| Central | 114 | Spring | Low | Drought | Y | 0.471 | -0.482 |
| Central | 115 | Fall | Low | Control | Y | 0.979 | - |
| Central | 115 | Spring | Low | Control | Y | 0.41 | - |
| Central | 116 | Spring | Low | Control | N | 0.001 | - |
| Central | 117 | Fall | Low | Drought | Y | 0.634 | -0.019 |
| Central | 117 | Spring | Low | Drought | Y | 0.052 | -1.439 |
| Central | 118 | Spring | Low | Drought | Y | 2.607 | 0.261 |
| Central | 119 | Fall | Low | Control | N | 0.484 | - |
| Central | 119 | Spring | Low | Control | Y | 1.483 | - |
| Central | 120 | Fall | Low | Drought | Y | 0.335 | -0.296 |
| Central | 120 | Spring | Low | Drought | Y | 1.227 | -0.066 |
| Southern | 1 | Fall | High | Drought | N | 0.001 | - |
| Southern | 1 | Spring | High | Drought | Y | 0.283 | -0.721 |
| Southern | 2 | Fall | High | Control | Y | 0.377 | 0.257 |
| Southern | 2 | Spring | High | Control | Y | 1.537 | 0.014 |
| Southern | 4 | Spring | High | Drought | Y | 0.61 | -0.387 |
| Southern | 5 | Fall | High | Warming | Y | 0.592 | 0.453 |
| Southern | 5 | Spring | High | Warming | Y | 0.15 | -0.996 |
| Southern | 7 | Fall | High | Drought | N | 0.001 | - |
| Southern | 7 | Spring | High | Drought | Y | 0.464 | -0.506 |
| Southern | 8 | Fall | High | Warming | Y | 0.148 | -0.149 |
| Southern | 8 | Spring | High | Warming | Y | 0.71 | -0.321 |
| Southern | 10 | Fall | High | Control | N | 0.001 | - |
| Southern | 10 | Spring | High | Control | N | 0.001 | - |
| Southern | 11 | Spring | High | Warming | Y | 0.89 | -0.223 |
| Southern | 13 | Spring | High | Control | Y | 0.673 | - |
| Southern | 14 | Spring | High | Warming | Y | 0.28 | -0.725 |
| Southern | 15 | Spring | High | Drought | Y | 0.513 | -0.462 |
| Southern | 16 | Spring | High | Control | Y | 2.582 | - |
| Southern | 17 | Fall | High | Warming | Y | 0.046 | -0.656 |
| Southern | 17 | Spring | High | Warming | Y | 0.47 | -0.5 |

| | | | | | | | |
|---|---|---|---|---|---|---|---|
| Southern | 18 | Fall | High | Drought | Y | 0.119 | -0.244 |
| Southern | 18 | Spring | High | Drought | Y | 0.641 | -0.365 |
| Southern | 19 | Fall | High | Control | Y | 0.04 | - |
| Southern | 19 | Spring | High | Control | Y | 1.156 | - |
| Southern | 101 | Fall | Low | Drought | Y | 0.753 | 0.161 |
| Southern | 101 | Spring | Low | Drought | Y | 0.598 | 0.176 |
| Southern | 102 | Fall | Low | Control | N | 0.001 | - |
| Southern | 102 | Spring | Low | Control | N | 0.001 | - |
| Southern | 103 | Fall | Low | Control | Y | 0.691 | - |
| Southern | 103 | Spring | Low | Control | Y | 0.389 | - |
| Southern | 104 | Spring | Low | Control | Y | 0.408 | 0 |
| Southern | 105 | Spring | Low | Drought | Y | 0.043 | -0.967 |
| Southern | 106 | Fall | Low | Drought | N | 0.001 | - |
| Southern | 106 | Spring | Low | Drought | Y | 0.325 | -0.089 |
| Southern | 107 | Spring | Low | Control | N | 0.001 | - |
| Southern | 108 | Fall | Low | Control | Y | 0.348 | -0.174 |
| Southern | 108 | Spring | Low | Control | N | 0.001 | - |
| Southern | 109 | Fall | Low | Drought | Y | 1.029 | 0.297 |
| Southern | 109 | Spring | Low | Drought | Y | 0.349 | -0.058 |
| Southern | 110 | Spring | Low | Drought | Y | 0.92 | 0.363 |

[a] The effect sizes were calculated where ANF activity was detected (i.e. 'Y' in column 'Fixation) and for both drought and warming samples.

**Table S4.** ANOVA results for metrics of plant diversity and cover. Tested with a Type II ANOVA.

| | **Rao's Q** | | **N fixing plant cover** | |
|---|---|---|---|---|
| | *F value* | *P* | *F value* | *P* |
| **Climate treatment** | 13.318 | **< 0.001** | 1.511 | 0.225 |
| **Season** | 0.051 | 0.822 | 0.119 | 0.731 |
| **Site** | 33.003 | **< 0.001** | 17.268 | **< 0.001** |
| **Diversity treatment** | 30.372 | **< 0.001** | 98.172 | **< 0.001** |
| **Diversity treatment x Site** | 25.641 | **< 0.001** | 11.512 | **< 0.001** |

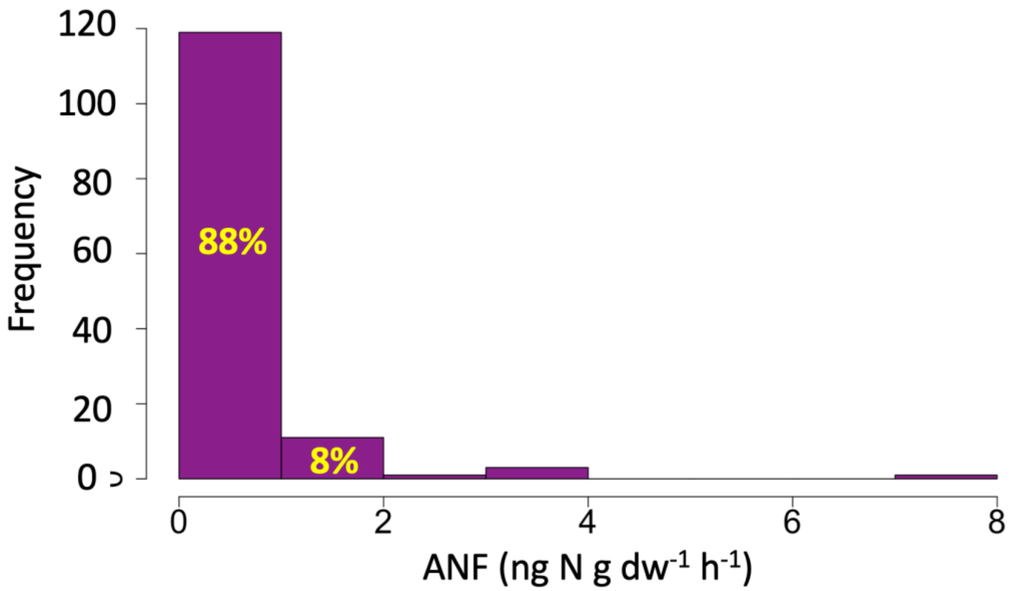


**Figure S2** Histogram of the calculated rates of asymbiotic nitrogen fixation (ANF, in ng N g dry weight$^{-1}$ h$^{-1}$) in soil sampled from high- and low-diversity plots during Fall 2018 sampling. Per cent values indicate values relative to total included in each bar. For instance, 88% of the ANF results fell between 0 and 1 ng N g dry weight$^{-1}$ h$^{-1}$.

**Table S5.** Soil biogeochemistry data across plots, sites, seasons, plant diversity and climate treatments.

| Site | Plot ID | Season | PlantDiv | Treatment | NH4-N (micg/g) | NO3-N (micg/g) | (Total Ni micg/g) | Total C (g/kg) | Total N (g/kg) | Total C/N (mass) | Available P (micg/g) | Available Fe (micg/g) | pH H2O | pH CaCl2 |
|---|---|---|---|---|---|---|---|---|---|---|---|---|---|---|
| | 41 | Spring | High | Control | 68.97 | 13.95 | 82.92 | 77.82 | 6.34 | 12.27 | 193.89 | 169.75 | 5.54 | 4.91 |
| | 42 | Spring | High | Warming | 54.71 | 16.67 | 71.38 | 81.01 | 6.00 | 13.50 | 45.29 | 121.97 | 5.72 | 4.83 |
| | 42 | Fall | High | Warming | 45.48 | 1.42 | 46.90 | 50.68 | 3.44 | 14.75 | 49.26 | 149.62 | 5.73 | 4.90 |
| | 44 | Spring | High | Drought | 76.43 | 7.29 | 83.72 | 69.76 | 5.18 | 13.47 | 73.35 | 179.30 | 5.67 | 4.76 |
| | 45 | Spring | High | Warming | 45.50 | 2.76 | 48.26 | 76.13 | 5.93 | 12.84 | 36.46 | 155.21 | 5.91 | 4.94 |
| | 47 | Spring | High | Drought | 38.60 | 1.34 | 39.93 | 75.11 | 5.83 | 12.88 | 155.05 | 207.55 | 5.60 | 4.64 |
| | 47 | Fall | High | Drought | 10.52 | 12.67 | 23.19 | 90.33 | 7.30 | 12.37 | 438.54 | 270.26 | 5.84 | 5.03 |
| | 48 | Spring | High | Control | 47.60 | 1.74 | 49.33 | 76.30 | 6.00 | 12.72 | 159.72 | 185.53 | 5.80 | 4.89 |
| | 48 | Fall | High | Control | 56.85 | 0.60 | 57.45 | 68.63 | 5.46 | 12.56 | 203.34 | 226.21 | 5.97 | 5.08 |
| | 49 | Spring | High | Drought | 54.41 | 1.11 | 55.52 | 76.07 | 6.16 | 12.35 | 76.47 | 186.78 | 5.70 | 4.83 |
| | 50 | Spring | High | Control | 38.82 | 5.49 | 44.32 | 67.98 | 5.35 | 12.72 | 44.77 | 166.84 | 5.90 | 4.88 |
| Northern | 50 | Fall | High | Control | 7.29 | 0.01 | 7.30 | 77.90 | 6.14 | 12.70 | 72.04 | 246.06 | 5.51 | 4.93 |
| | 51 | Spring | High | Warming | 58.09 | 5.60 | 63.69 | 76.23 | 5.59 | 13.64 | 86.86 | 178.06 | 5.70 | 4.88 |
| | 51 | Fall | High | Warming | 19.67 | 2.80 | 22.47 | 93.58 | 6.68 | 14.02 | 79.08 | 208.10 | 5.90 | 5.13 |
| | 53 | Spring | High | Control | 44.61 | 1.88 | 46.49 | 71.65 | 5.81 | 12.33 | 76.47 | 202.15 | 5.78 | 4.78 |
| | 54 | Spring | High | Warming | 76.85 | 3.57 | 80.42 | 68.41 | 5.05 | 13.56 | 87.90 | 216.69 | 5.67 | 4.80 |
| | 56 | Fall | High | Drought | 13.53 | 0.13 | 13.66 | 55.64 | 3.98 | 13.98 | 55.84 | 372.97 | 5.74 | 4.96 |
| | 56 | Spring | High | Drought | 60.23 | 2.61 | 62.84 | 81.84 | 6.44 | 12.72 | 51.53 | 140.67 | 5.98 | 4.97 |
| | 58 | Fall | High | Drought | 21.07 | 4.41 | 25.48 | 62.22 | 4.63 | 13.44 | 37.96 | 193.01 | 5.81 | 4.91 |
| | 58 | Spring | High | Drought | 32.63 | 3.47 | 36.10 | 61.29 | 4.31 | 14.21 | 19.32 | 136.10 | 5.81 | 4.77 |
| | 59 | Fall | High | Control | 81.48 | 0.47 | 81.95 | 84.91 | 7.05 | 12.05 | 250.84 | 300.72 | 6.31 | 5.26 |
| | 59 | Spring | High | Control | 52.99 | 1.79 | 54.79 | 74.06 | 5.73 | 12.92 | 63.48 | 158.95 | 5.82 | 4.80 |
| | 60 | Fall | High | Warming | 59.02 | 1.83 | 60.85 | 67.38 | 5.20 | 12.95 | 53.92 | 228.18 | 5.60 | 4.69 |
| | 60 | Spring | High | Warming | 62.65 | 0.01 | 62.66 | 70.10 | 5.01 | 13.98 | 28.67 | 195.92 | 5.74 | 4.59 |
| | 121 | Spring | Low | Control | 59.36 | 5.23 | 64.59 | 34.02 | 2.86 | 11.88 | 162.84 | 185.12 | 5.74 | 4.93 |
| | 121 | Fall | Low | Control | 34.36 | 1.02 | 35.38 | 69.67 | 5.67 | 12.30 | 141.58 | 243.59 | 5.59 | 4.72 |
| | 122 | Spring | Low | Drought | 202.47 | 5.58 | 208.05 | 48.36 | 3.37 | 14.35 | 67.81 | 188.03 | 5.77 | 4.75 |
| | 122 | Fall | Low | Drought | 29.67 | 0.01 | 29.68 | 68.10 | 5.29 | 12.88 | 140.00 | 209.68 | 5.96 | 4.91 |
| | 123 | Spring | Low | Control | 39.31 | 6.72 | 46.03 | 82.27 | 6.95 | 11.85 | 43.22 | 179.72 | 5.86 | 4.79 |
| | 123 | Fall | Low | Control | 24.49 | 0.01 | 24.50 | 76.67 | 6.48 | 11.84 | 42.39 | 239.64 | 5.90 | 5.09 |
| | 124 | Spring | Low | Drought | 147.16 | 4.35 | 151.52 | 68.70 | 5.60 | 12.26 | 135.18 | 168.50 | 6.02 | 5.18 |
| | 125 | Spring | Low | Control | 75.32 | 0.01 | 75.33 | 73.40 | 5.78 | 12.70 | 102.96 | 186.78 | 5.68 | 4.77 |
| Northern | 126 | Spring | Low | Drought | 56.74 | 0.84 | 57.57 | 77.28 | 6.26 | 12.35 | 94.29 | 195.92 | 5.92 | 4.98 |
| | 126 | Fall | Low | Drought | 3.62 | 0.01 | 3.63 | 114.18 | 8.64 | 13.21 | 114.46 | 237.19 | 5.65 | 4.88 |
| | 127 | Spring | Low | Drought | 71.89 | 0.87 | 72.76 | 87.04 | 7.44 | 11.70 | 188.69 | 101.20 | 5.74 | 4.98 |
| | 127 | Fall | Low | Drought | 8.31 | 0.06 | 8.37 | 82.65 | 6.38 | 12.96 | 98.86 | 182.67 | 6.00 | 4.91 |
| | 128 | Spring | Low | Control | 37.59 | 21.57 | 59.17 | 94.31 | 8.05 | 11.72 | 212.69 | 231.65 | 5.60 | 4.70 |
| | 128 | Fall | Low | Control | 1.63 | 0.01 | 1.64 | 97.30 | 8.52 | 11.43 | 332.40 | 313.17 | 6.21 | 5.16 |
| | 129 | Spring | Low | Control | 67.23 | 0.01 | 67.24 | 67.73 | 5.50 | 12.31 | 109.72 | 175.98 | 5.83 | 4.88 |
| | 130 | Spring | Low | Drought | 109.26 | 1.73 | 110.99 | 79.37 | 5.81 | 13.67 | 37.50 | 164.35 | 5.71 | 4.96 |
| | 21 | Fall | High | Drought | 40.18 | 4.94 | 45.12 | 107.70 | 8.00 | 13.46 | 15.89 | 53.19 | 5.70 | 4.83 |
| Central | 21 | Spring | High | Drought | 18.12 | 3.06 | 21.19 | 91.13 | 7.43 | 12.27 | 17.80 | 72.24 | 5.71 | 4.96 |
| | 22 | Fall | High | Control | 30.45 | 4.32 | 34.77 | 95.00 | 7.10 | 13.38 | 6.85 | 54.36 | 5.46 | 4.66 |

| | | | | | | | | | | | | | | |
|---|---|---|---|---|---|---|---|---|---|---|---|---|---|---|
| | 22 | Spring | High | Control | 19.21 | 7.71 | 26.91 | 113.80 | 8.62 | 13.21 | 21.40 | 63.16 | 5.68 | 4.65 |
| | 23 | Fall | High | Warming | 42.86 | 14.39 | 57.25 | 68.70 | 5.40 | 12.72 | 6.10 | 73.13 | 5.25 | 4.77 |
| | 23 | Spring | High | Warming | 7.98 | 2.71 | 10.69 | 66.84 | 5.41 | 12.36 | 12.74 | 52.10 | 5.67 | 4.87 |
| | 25 | Fall | High | Drought | 40.77 | 2.13 | 42.90 | 80.50 | 6.10 | 13.20 | 5.65 | 75.95 | 5.58 | 4.87 |
| | 25 | Spring | High | Drought | 14.33 | 0.01 | 14.34 | 66.83 | 5.41 | 12.35 | 11.69 | 62.31 | 5.71 | 4.70 |
| | 26 | Spring | High | Warming | 20.35 | 0.71 | 21.06 | 102.46 | 8.35 | 12.27 | 14.40 | 63.03 | 5.59 | 4.81 |
| | 28 | Spring | High | Drought | 14.54 | 1.23 | 15.76 | 105.48 | 8.37 | 12.61 | 9.82 | 58.90 | 5.92 | 4.93 |
| | 29 | Fall | High | Control | 70.17 | 1.80 | 71.97 | 64.20 | 5.20 | 12.35 | 6.32 | 80.11 | 5.62 | 4.80 |
| | 29 | Spring | High | Control | 16.45 | 0.54 | 16.99 | 70.61 | 5.88 | 12.01 | 13.85 | 77.84 | 5.57 | 4.74 |
| | 30 | Fall | High | Drought | 56.89 | 3.59 | 60.48 | 91.30 | 7.20 | 12.68 | 48.75 | 77.94 | 5.52 | 4.86 |
| | 30 | Spring | High | Drought | 16.17 | 1.05 | 17.22 | 109.27 | 8.31 | 13.16 | 11.60 | 60.26 | 5.36 | 4.72 |
| | 31 | Spring | High | Control | 6.88 | 2.69 | 9.57 | 97.83 | 7.59 | 12.88 | 12.14 | 64.94 | 5.74 | 4.79 |
| | 32 | Fall | High | Warming | 27.91 | 1.51 | 29.42 | 72.10 | 5.60 | 12.88 | 5.63 | 74.37 | 5.60 | 4.85 |
| | 32 | Spring | High | Warming | 16.29 | 2.70 | 18.99 | 89.06 | 7.00 | 12.73 | 16.42 | 65.93 | 5.82 | 4.88 |
| | 34 | Fall | High | Control | 51.82 | 2.46 | 54.28 | 97.20 | 7.10 | 13.69 | 7.71 | 79.70 | 5.75 | 5.00 |
| | 34 | Spring | High | Control | 14.90 | 2.38 | 17.28 | 98.45 | 7.35 | 13.40 | 17.55 | 66.24 | 5.76 | 4.86 |
| | 35 | Fall | High | Warming | 83.26 | 6.35 | 89.61 | 93.00 | 7.10 | 13.10 | 6.03 | 68.67 | 5.78 | 5.00 |
| | 35 | Spring | High | Warming | 19.88 | 1.62 | 21.50 | 98.55 | 7.55 | 13.05 | 18.73 | 64.00 | 5.94 | 5.15 |
| | 38 | Spring | High | Drought | 14.34 | 0.94 | 15.28 | 85.42 | 6.58 | 12.99 | 15.05 | 60.17 | 5.88 | 5.03 |
| | 39 | Spring | High | Control | 21.85 | 0.44 | 22.29 | 68.40 | 5.40 | 12.67 | 7.67 | 55.78 | 5.79 | 4.93 |
| | 40 | Spring | High | Warming | 17.82 | 6.37 | 24.19 | 97.80 | 7.33 | 13.34 | 8.40 | 65.27 | 5.79 | 4.93 |
| | 111 | Fall | Low | Control | 78.63 | 6.93 | 85.56 | 109.50 | 7.90 | 13.86 | 7.63 | 81.47 | 5.74 | 5.16 |
| | 111 | Spring | Low | Control | 14.43 | 1.03 | 15.46 | 113.56 | 8.38 | 13.56 | 22.05 | 71.20 | 5.60 | 5.18 |
| | 112 | Fall | Low | Drought | 45.02 | 5.78 | 50.80 | 104.10 | 5.60 | 18.59 | 4.98 | 68.93 | 5.76 | 5.23 |
| | 112 | Spring | Low | Drought | 13.74 | 9.42 | 23.16 | 101.15 | 7.63 | 13.25 | 26.05 | 59.69 | 5.73 | 5.03 |
| | 113 | Spring | Low | Control | 18.37 | 0.86 | 19.24 | 103.39 | 7.31 | 14.15 | 10.48 | 69.04 | 5.93 | 5.21 |
| | 114 | Spring | Low | Drought | 13.40 | 1.35 | 14.74 | 95.70 | 6.89 | 13.89 | 22.60 | 67.10 | 6.02 | 5.12 |
| | 115 | Fall | Low | Control | 87.20 | 7.00 | 94.20 | 100.50 | 7.40 | 13.58 | 50.18 | 78.79 | 5.99 | 5.26 |
| Central | 115 | Spring | Low | Control | 15.25 | 4.71 | 19.96 | 122.36 | 8.96 | 13.66 | 17.35 | 63.32 | 5.87 | 5.13 |
| | 116 | Spring | Low | Control | 12.19 | 0.14 | 12.33 | 107.37 | 7.48 | 14.36 | 12.52 | 74.57 | 5.75 | 4.94 |
| | 117 | Fall | Low | Drought | 81.21 | 40.42 | 121.63 | 107.80 | 7.90 | 13.65 | 13.90 | 86.61 | 5.94 | 5.57 |
| | 117 | Spring | Low | Drought | 9.35 | 3.46 | 12.81 | 106.97 | 7.51 | 14.24 | 15.33 | 79.35 | 6.21 | 5.55 |
| | 118 | Spring | Low | Drought | 10.81 | 4.01 | 14.82 | 91.24 | 6.74 | 13.54 | 14.79 | 65.22 | 6.06 | 5.12 |
| | 119 | Fall | Low | Control | 43.79 | 1.63 | 45.42 | 57.60 | 4.30 | 13.40 | 4.98 | 83.02 | 5.91 | 5.16 |
| | 119 | Spring | Low | Control | 9.78 | 0.89 | 10.67 | 93.43 | 6.45 | 14.50 | 10.03 | 57.90 | 6.03 | 5.16 |
| | 120 | Fall | Low | Drought | 32.51 | 6.23 | 38.74 | 68.40 | 5.00 | 13.68 | 7.49 | 87.13 | 5.59 | 5.05 |
| | 120 | Spring | Low | Drought | 20.54 | 7.90 | 28.44 | 109.54 | 7.19 | 15.23 | 29.92 | 80.76 | 6.08 | 5.27 |
| | 1 | Fall | High | Drought | 17.00 | 1.94 | 18.94 | 59.77 | 5.06 | 11.80 | 76.99 | 211.72 | 6.47 | 5.78 |
| | 1 | Spring | High | Drought | 19.25 | 1.85 | 21.10 | 40.95 | 3.69 | 11.09 | 44.07 | 202.84 | 6.26 | 5.49 |
| | 2 | Fall | High | Control | 17.94 | 1.33 | 19.27 | 34.65 | 3.15 | 11.00 | 20.75 | 198.60 | 6.31 | 5.65 |
| | 2 | Spring | High | Control | 26.02 | 2.42 | 28.45 | 42.25 | 3.95 | 10.71 | 63.89 | 193.71 | 6.23 | 5.38 |
| | 4 | Spring | High | Drought | 71.37 | 0.01 | 71.38 | 31.55 | 2.92 | 10.80 | 43.47 | 144.91 | 6.33 | 5.46 |
| | 5 | Fall | High | Warming | 7.83 | 1.52 | 9.35 | 30.51 | 2.96 | 10.31 | 20.28 | 148.69 | 6.20 | 5.56 |
| Southern | 5 | Spring | High | Warming | 26.55 | 2.04 | 28.59 | 33.35 | 3.24 | 10.30 | 19.97 | 144.38 | 5.56 | 5.24 |
| | 7 | Fall | High | Drought | 12.58 | 2.40 | 14.98 | 26.72 | 2.39 | 11.19 | 69.51 | 175.01 | 6.69 | 5.95 |
| | 7 | Spring | High | Drought | 15.32 | 7.77 | 23.09 | 26.50 | 2.40 | 11.05 | 33.08 | 174.14 | 6.42 | 5.53 |
| | 8 | Fall | High | Warming | 30.36 | 2.54 | 32.90 | 47.61 | 4.29 | 11.10 | 56.90 | 173.70 | 6.43 | 5.80 |
| | 8 | Spring | High | Warming | 17.85 | 2.84 | 20.69 | 26.73 | 2.46 | 10.85 | 29.86 | 170.63 | 6.38 | 5.47 |
| | 10 | Fall | High | Control | 9.42 | 2.73 | 12.15 | 51.13 | 4.77 | 10.71 | 46.41 | 176.32 | 6.09 | 5.41 |
| | 10 | Spring | High | Control | 20.45 | 12.87 | 33.32 | 38.37 | 3.58 | 10.73 | 42.23 | 173.95 | 6.19 | 5.36 |

| | | | | | | | | | | | | | | |
|---|---|---|---|---|---|---|---|---|---|---|---|---|---|---|
| Southern | 11 | Spring | High | Warming | 23.07 | 7.95 | 31.02 | 38.77 | 3.48 | 11.14 | 41.68 | 168.15 | 6.45 | 5.65 |
| | 13 | Spring | High | Control | 47.08 | 4.67 | 51.75 | 31.47 | 2.88 | 10.93 | 29.05 | 165.92 | 6.26 | 5.41 |
| | 14 | Spring | High | Warming | 41.45 | 6.58 | 48.03 | 27.62 | 2.53 | 10.91 | 14.11 | 155.25 | 6.37 | 5.55 |
| | 15 | Spring | High | Drought | 14.15 | 0.01 | 14.16 | 45.10 | 4.22 | 10.68 | 41.09 | 193.65 | 6.16 | 5.32 |
| | 16 | Spring | High | Control | 17.26 | 3.18 | 20.43 | 38.94 | 3.54 | 10.99 | 37.45 | 167.42 | 6.24 | 5.46 |
| | 17 | Fall | High | Warming | 0.37 | 2.26 | 2.63 | 34.13 | 3.26 | 10.48 | 58.37 | 168.87 | 6.34 | 5.66 |
| | 17 | Spring | High | Warming | 22.14 | 2.20 | 24.34 | 26.12 | 2.54 | 10.27 | 76.25 | 171.52 | 6.33 | 5.56 |
| | 18 | Fall | High | Drought | 12.46 | 0.90 | 13.36 | 34.36 | 3.20 | 10.75 | 45.40 | 170.74 | 6.44 | 5.74 |
| | 18 | Spring | High | Drought | 28.63 | 2.69 | 31.32 | 32.95 | 3.04 | 10.85 | 34.45 | 126.94 | 6.42 | 5.56 |
| | 19 | Fall | High | Control | 18.95 | 0.94 | 19.89 | 40.19 | 3.65 | 11.00 | 45.92 | 166.34 | 6.40 | 5.79 |
| | 19 | Spring | High | Control | 32.76 | 1.06 | 33.82 | 40.04 | 3.66 | 10.95 | 43.24 | 185.87 | 6.34 | 5.42 |
| | 101 | Fall | Low | Drought | 14.36 | 6.39 | 20.75 | 70.37 | 5.91 | 11.91 | 60.45 | 210.38 | 6.55 | 5.85 |
| | 101 | Spring | Low | Drought | 41.92 | 4.71 | 46.63 | 34.71 | 3.05 | 11.39 | 49.08 | 228.22 | 6.31 | 5.67 |
| | 102 | Fall | Low | Control | 35.81 | 1.75 | 37.56 | 45.60 | 4.07 | 11.21 | 61.00 | 199.08 | 6.70 | 6.03 |
| | 102 | Spring | Low | Control | 75.57 | 1.44 | 77.01 | 41.07 | 3.58 | 11.46 | 23.97 | 175.66 | 6.32 | 5.54 |
| | 103 | Fall | Low | Control | 15.69 | 0.99 | 16.68 | 50.96 | 4.54 | 11.22 | 52.86 | 159.34 | 6.67 | 6.15 |
| | 103 | Spring | Low | Control | 37.08 | 4.73 | 41.80 | 35.40 | 3.18 | 11.12 | 19.24 | 176.87 | 6.26 | 5.50 |
| | 104 | Spring | Low | Control | 51.37 | 3.31 | 54.68 | 46.98 | 4.28 | 10.97 | 55.22 | 196.36 | 6.37 | 5.65 |
| | 105 | Spring | Low | Drought | 65.17 | 2.47 | 67.64 | 52.37 | 4.41 | 11.88 | 31.98 | 179.56 | 6.40 | 5.73 |
| | 106 | Fall | Low | Drought | 26.77 | 0.01 | 26.78 | 58.92 | 4.79 | 12.31 | 41.70 | 163.62 | 6.59 | 5.94 |
| | 106 | Spring | Low | Drought | 50.16 | 1.13 | 51.29 | 32.25 | 2.92 | 11.04 | 51.40 | 167.01 | 6.46 | 5.71 |
| | 107 | Spring | Low | Control | 53.91 | 0.01 | 53.92 | 25.37 | 2.28 | 11.15 | 27.96 | 129.65 | 6.49 | 5.73 |
| | 108 | Fall | Low | Control | 15.12 | 1.18 | 16.30 | 53.28 | 4.75 | 11.21 | 55.53 | 181.50 | 6.56 | 5.92 |
| | 108 | Spring | Low | Control | 54.54 | 0.01 | 54.55 | 49.68 | 4.39 | 11.32 | 64.41 | 188.93 | 6.38 | 5.71 |
| | 109 | Fall | Low | Drought | 21.81 | 2.42 | 24.23 | 41.10 | 3.97 | 10.35 | 9.63 | 155.96 | 6.33 | 6.14 |
| | 109 | Spring | Low | Drought | 32.06 | 0.01 | 32.07 | 38.14 | 3.64 | 10.49 | 21.40 | 151.82 | 6.67 | 5.92 |
| | 110 | Spring | Low | Drought | 53.96 | 5.45 | 59.41 | 44.49 | 4.05 | 10.99 | 43.09 | 174.52 | 6.42 | 5.69 |
